# Citrus vascular proteomics highlights the role of peroxidases and serine proteases during Huanglongbing disease progression

**DOI:** 10.1101/2020.04.05.025718

**Authors:** Jessica Y. Franco, Shree P. Thapa, Zhiqian Pang, Fatta B. Gurung, Thomas W.H. Liebrand, Danielle M Stevens, Veronica Ancona, Nian Wang, Gitta Coaker

## Abstract

Huanglongbing (HLB) is the most devastating and widespread citrus disease. All commercial citrus varieties are susceptible to the HLB-associated bacterium, *Candidatus* Liberibacter asiaticus (*C*Las), which resides in the phloem. The phloem is part of the plant vascular system and is involved in sugar transport. To investigate the plant response to *C*Las, we enriched for proteins surrounding the phloem in an HLB susceptible sweet orange variety, Washington navel (*Citrus sinensis* (L) Osbeck). Quantitative proteomics revealed global changes in the citrus proteome after *C*Las inoculation. Plant metabolism and translation were suppressed, while defense-related proteins such as peroxidases, proteases and protease inhibitors were induced in the vasculature. Transcript accumulation and enzymatic activity of plant peroxidases in CLas infected sweet orange varieties under greenhouse and field conditions were assessed. While peroxidase transcript accumulation was induced in *C*Las infected sweet orange varieties, peroxidase enzymatic activity varied. Specific serine proteases were upregulated in Washington navel in the presence of *C*Las based on quantitative proteomics. Subsequent activity-based protein profiling revealed increased activity of two serine proteases, and reduced activity of one protease in two *C. sinensis* sweet orange varieties under greenhouse and field conditions. The observations in the current study highlight global reprogramming of the citrus vascular proteome and differential regulation of enzyme classes in response to *C*Las infection. These results open an avenue for further investigation of diverse responses to HLB across different environmental conditions and citrus genotypes.

## Introduction

Currently, the citrus industry is facing a major threat from the bacterial vector-borne disease huanglongbing (HLB), also known as citrus greening. The HLB-associated pathogen, *Candidatus* Liberibacter asiaticus (*C*Las), is spread by *Diaphorina citri*, a hemipteran insect. All commercial citrus varieties are susceptible to *C*Las and current interventions such as *D. citri* control and infected tree removal have had little success in controlling the disease (1). Sweet orange varieties (*Citrus sinensis*) exhibit the fastest decline, while lemon (*C. latifolia*), mandarins (*C. reticulata*), and tangerines (*C. reticulata*) present delayed symptom progression but ultimately succumb to HLB (2, 3). In the United States, HLB was first diagnosed in 2005 in Florida and has since spread to Texas, Louisiana, South Carolina, Georgia and California (4). In California, *C*Las positive citrus was detected in 2012 and there are currently 1,788 positive plants identified by qPCR (5, 6). In Florida, it is estimated that 95% of mature trees in commercial groves are infected, resulting in a loss of over 7.8 billion dollars in revenue and 7,500 jobs since 2007 (7). For decades, Florida has been the primary citrus producer in the United States. However, due to losses from HLB, California is now the primary citrus producing area (8).

Plant pathogens can be directly deposited into the vascular system (xylem or phloem) by piercing sucking insects. *D. citri* deposits *C*Las into plant phloem sieve elements, where the bacteria proliferate (9). The phloem is rich in sugars, amino acids, and proteins (10). *C*Las infected leaf tissues exhibit increased deposition of callose, a beta 1-3 glucan, and accumulation of starch that constricts symplastic transport, which contributes to symptom development (11, 12). HLB symptoms are characterized by yellowing of the shoots and leaves as well as hard, misshapen, and bitter fruit (13, 14). Early detection efforts have been challenging to develop due to the long latent period and the uneven distribution of *C*Las within its plant host (15). The inability to culture *C*Las has hampered progress in studying the pathogen and its virulence mechanisms.

One of the paramount mechanisms of bacterial pathogenesis is the presence of secretion systems facilitating the translocation of proteins, or effectors, to the plant host (16). *C*Las possesses the type I and general sec secretion system (17, 18). The sec pathway is critical for virulence in other phloem-colonizing pathogens including mollicutes (*Ca*. Phytoplasma) (19). Most *C*Las sec-dependent effectors (SDEs) were highly expressed *in planta*, indicating they are important for pathogen proliferation (18, 20, 21). Transient expression of a conserved *C*Las SDE, SDE1, induces cell death, starch accumulation, and callose deposition in *Nicotiana benthamiana* (22). In citrus, SDE1 inhibits the activity of immune related proteases which may serve as a strategy to dampen defense responses (23). Multiple secreted *C*Las proteins can suppress immune responses (23–26). Upon pathogen perception, the production of reactive oxygen species (ROS), including hydrogen peroxide, serve as an antimicrobial as well as a secondary immune signal (27, 28). *C*Las peroxiredoxins, LasBCP and LasdPrx5, increased oxidative stress tolerance in *Liberibacter crescens* (a culturable surrogate for *C*Las) by suppressing H_2_O_2_ accumulation (24, 29). *C*Las encodes a salicylic acid hydroxylase that degrades salicylic acid (SA), a plant defense hormone (26). Collectively, these data demonstrate that *C*Las actively manipulates plant defense responses.

In order to understand the *C. sinensis* response to *C*Las, previous studies have investigated changes in gene and protein expression during infection by microarray, RNAseq, and proteomics (30–32). Microarray analyses revealed induction of genes associated with sugar metabolism, plant defenses, phytohormones, and cell wall metabolism in infected leaf samples (33). Transcripts encoding proteins involved in sugar and starch metabolism, stress responses, transport, detoxification, and lipid metabolism were significantly upregulated during *C*Las infection, while processes involved in photosynthesis were downregulated in mature leaf, stem, and root tissues (30, 31, 34, 35). *C*Las induces plant defense responses in leaf tissue, including the transcription of pathogenesis-related (PR) genes, chitinases, callose synthases, and defense-related WRKY transcription factors (11, 31, 33, 35, 36). Consistent with these observations, infected *C. sinensis* midveins exhibit callose and starch accumulation in the sieve tubes that lead to phloem plugging and collapse (33, 34, 37). However, in roots, genes encoding callose hydrolases were induced and callose synthases were repressed (35). These data illustrate that not all tissue types respond ubiquitously and highlight tissue-specific responses during *C*Las infection.

Multiple studies have elucidated host responses in intact leaf, root, and fruit samples (30, 31, 35, 36). However, *C*Las is limited to phloem sieve elements and there is a lack of proteomic studies on vascular exudates. In this study, we performed comparative proteomic analyses to investigate changes in the vascular proteome after *C*Las infection in the sweet orange variety Washington Navel (*Citrus sinensis* (L) Osbeck). Our results highlight the accumulation of peroxidases and proteases upon *C*Las infection, indicating the activation of plant defense responses. Multiple citrus peroxidases were induced at the transcript and protein level during infection across different citrus genotypes and environmental conditions. Serine protease activity in leaf tissues also dynamically changed after *C*Las infection under greenhouse and field conditions. This study advances our understanding of vascular changes and highlights the important role of peroxidases and proteases during *C*Las infection.

## Experimental Procedures

### Experimental Design and Statistical Rationale

For vascular proteomics, eight Washington Naval trees grafted onto Carrizo rootstock were grown in the UC Davis Contained Research Facility, a Biosafety level 3 greenhouse. Seven months after grafting onto Carrizo rootstock, plants were graft-inoculated or mock-inoculated with the *C*Las isolate, HHCA obtained from Hacienda Heights, California, USA. Plants were grown in a completely randomized design, tested for *C*Las titers by qPCR monthly, and vascular exudates were harvested ten months post graft-inoculation. Vascular exudates from four mock and four *C*Las-graft inoculated Washington navel trees were extracted using the centrifugation method (10). For mass spectrometry analyses, samples were blinded to the operator. To quantify differentially changing proteins, label-free quantification using MaxQuant (v 1.5.1.0) was performed. Protein identification search parameters included a decoy search database in addition to a False Discovery Rate (FDR) of < 0.05. Statistical analyses were performed on four individual trees per treatment (n=4). Differentially changing proteins were identified using a two-sample student’s t-test with an alpha-value of 0.05 for truncation.

### Plant Growth and Inoculation Conditions

#### Washington Navel

Proteomic analyses were performed on citrus grown and maintained at the University of California, Davis Contained Research Facility greenhouse. The greenhouse temperature was maintained at 27°C. High pressure sodium lights were provided as supplemental lighting for 16 hours. The greenhouse was covered with shade cloth (50% shade) between mid-April and mid-October. Eight Washington navel (*Citrus sinensis* (L.) Osbeck) buds were grafted onto Carrizo rootstock using the inverted T budding method. After seven months, four Washington navel scions were mock or *C*Las graft-inoculated on February 2014. Samples were harvested 10 months post-graft inoculation and tested for *C*Las presence by qPCR each month. A random sampling of 6-8 leaves was taken from each tree and pooled for DNA extraction. DNA was extracted using the Qiagen MagAttract plant DNA extraction kit (Qiagen, Valencia, CA, USA). *C*Las presence was detected by qPCR using the USDA-APHIS-PPQ protocol (38). *C*Las primers amplifying the 16s rDNA (HLBas/HLBr) were used for *C*Las detection. The citrus cytochrome oxidase (COXf/r) primers and probe were used as an internal control (38). All primers, gene names, and accession numbers are provided in Supplementary Table S1.

#### Hamlin sweet orange

Hamlin sweet oranges (*Citrus sinensis* (L) Osbeck) were grown in the greenhouse at 25°C. Trees were exposed to *C*Las positive *D. citri* for 2 months. Psyllids were removed and plants were transferred to a psyllid-free greenhouse and sampled one year after *C*Las transmission. *C*Las detection was performed by qPCR using primers that amplify the β-operon (39). Samples were collected from uninfected and symptomatic Hamlin sweet orange, flash frozen, and sent to the Contained Research Facility on dry ice for further processing.

#### N-33

Mature N-33 (*Citrus sinensis* (L) Osbeck) trees were grown in a commercial orchard in Donna, TX. Leaf samples from symptomatic trees grown in the field and uninfected trees from an adjacent screenhouse were collected and flash frozen in liquid nitrogen. *C*Las detection was performed using qPCR with primers that amplify the 16s rDNA (HLBas/HLBr) and the *C. sinensis* GAPDH gene was used as an internal control. Samples were sent on dry ice to the Contained Research Facility at the University of California, Davis for further processing.

### Protein extraction for Mass Spectrometry

Citrus vascular exudates were enriched by centrifugation (10). Stems were collected from 17-month-old greenhouse plants and the outer bark was manually removed. The inner part of the bark was briefly rinsed with deionized water and dried with Kimwipes. The bark was cut into 1 cm pieces using a razor blade and placed into a 0.65 mL Eppendorf tube with a small hole in the bottom. This tube was then placed into a 1.7 mL Eppendorf collection tube and centrifuged at 12,000 x g for 15 min at 25°C. Vascular extracts were flash frozen and stored at −80°C. Samples were resuspended in 2x Laemmli buffer and quantified for protein concentration using the Pierce 660nm Protein Assay Reagent (Thermo Fisher Scientific).

A total of 85 μg of protein per sample was subjected to one-dimensional SDS-PAGE using an 8-16% precise protein gradient gel (Thermo Fisher Scientific). Proteins were separated for 1 cm in the resolving gel and the entire lane was excised and cut into equal pieces. In-gel reduction was performed by adding 10 mM dithiothreitol in 50 mM ammonium bicarbonate for 30 min at 56°C. Alkylation was performed by adding 55 mM iodoacetamide in 50 mM of ammonium bicarbonate for 20 min in the dark with shaking at room temperature. In-gel tryptic digestion was performed using 300 ng of trypsin in 50 mM ammonium bicarbonate (40). The digested peptides were dried in a vacuum concentrator and then solubilized in 120 μL of 2% acetonitrile, 0.1% trifluoroacetic acid for LC-MS/MS analysis.

### LC-MS/MS

Peptides were submitted to the Genomics Center Proteomics Core at the University of California, Davis for liquid chromatography tandem mass spectrometry (LC)-MS/MS. Peptides were analyzed by MS as described previously (23). The LC-MS/MS system configuration consisted of a CTC Pal autosampler (LEAP Technologies) and Paradigm HPLC device (Michrom BioResources) coupled to a QExactive hybrid quadrupole Orbitrap mass spectrometer (Thermo Fisher Scientific) with a CaptiveSpray ionization source (Michrom BioResources). Peptides were reconstituted in 2% acetonitrile and 0.1% formic acid and were washed on a Michrom C18 trap then were eluted and separated on a Michrom Magic C18AQ (200 μm × 150 mm) capillary reverse-phase column at a flow rate of 3 μL/min. A 120 min gradient was applied with a 2% to 35% B (100% acetonitrile) for 100 min, a 35% B to 80% B for 7 min and 80% B for 2 min. Then a decrease of 80% to 5% B in 1 min followed by 98% A (0.1% formic acid) for 10 min. The QExactive was operated in Data-Dependent Acquisition (DDA) mode with a top-15 method. Spray voltage was set to 2.2 kV. The scan range was set to 350–1600 m/z, the maximum injection time was 30 ms and automatic gain control was set to 1 × 10^6^. Precursor resolution was set to 70,000. For MS/MS, the maximum injection time was 50 ms, the isolation window was 1.6 m/z, the scan range 200–2000 m/z, automatic gain control was set to 5 × 104 and normalized collision energy was 27%. The dynamic exclusion window was set to 15 s and fragment product resolution was 17,500. An intensity threshold of 1 × 104 was applied and the underfill ratio was 1%.

### Peptide identification, analyses and quantification

The raw MS/MS data files were imported into MaxQuant v1.5.1.0 for label-free intensity-based quantification (41). The database search engine Andromeda (42) was used to search MS/MS spectra against the a total of 81192 protein sequences from *C. clementina* (v1.0) and *C. sinensis* (v1.0) databases downloaded from Phytozome (43) and the *Candidatus* Liberibacter asiaticus (*C*Las) proteome database (44) downloaded from Uniprot with a tolerance level of 20 ppm for the first search and 6 ppm for the main search. Trypsin/P was set as the enzyme and two missed cleavages were allowed. Carbamidomethyl (Cys) was set as a fixed modification. Protein N-terminal acetylation and Methionine (Met) oxidation were set as variable modifications. The maximum number of modifications per peptide was set as five and the predefined contaminants from MaxQuant were included in the search database. The ‘match between runs’ feature was checked with a match time window of 0.7 min and an alignment time window of 20 min. The FDR for protein level and peptide spectrum match was set to 1%. The minimum peptide length was 6, minimum razor was changed to 0, and minimum unique peptides was set to 1. The minimum ratio count for protein quantification was set to 2. The other MaxQuant settings were default. The total peptide intensities for each replicate were summed and a normalization factor was calculated for each sample (41). This normalization factor was applied based on the least overall proteome change. Peptide ratios between samples were then calculated to obtain a pairwise protein ratio matrix between samples, which was subsequently used to rescale the cumulative intensity in each sample and provides the label-free intensity (LFQ) value (41). Parameters used in MaxQuant can be found in Supplementary Table S2. The MaxQuant output file “Protein Groups” was imported into Perseus 1.5.5.3 (45). Contaminants, reverse hits, and proteins identified only by modified peptides were excluded in subsequent analysis. The LFQ intensities were log_2_-transformed. Proteins not consistently identified in at least two out of the four replicates in at least one group were discarded. Missing values were imputed with values from a normal distribution of the obtained intensities using default settings (width 0.5, downshift 1.8). Differentially changing proteins were identified using a two-tailed Student’s t-test, p<0.05, with a permutation-based FDR of 0.05 to correct for multiple testing. The mass spectrometry proteomics data have been deposited to the ProteomeXchange Consortium via the PRIDE (46) partner repository with the dataset identifier PXD020703. The PRIDE reviewer login is: Username, Password: e3qL8mhM. Peptide identifications can be accessed by MS-Viewer (47) search key, 9mv75ckfbb.

### Comparative analyses of proteomics data

Initial data visualization was performed using Perseus (Version 1.5.5.3) software. The heatmap was generated by hierarchical clusters for both individual protein expression and replication was performed using Euclidean distance (48). Linkage was calculated using preprocessed k-means with 300 clusters, 10 maximal iterations, and 1 restart. Principle component analyses was performed using untransformed label-free values.

### Comparison of vascular proteomic studies

Homologs of citrus proteins identified by mass spectrometry were identified using BLAST analyses (version 2.2.18) against *Arabidopsis thaliana* (49, 50) and *Cucumis sativus* (51) phloem proteins with an E-value of 1e-10 required for protein identification. The Venn diagram was generated using the vennueler R package (52).

Gene ontology analyses were performed by using Phytozome (43). All identified protein groups were used as background. Gene ontology enrichment for differentially changing proteins using the Bonferroni test was used for correction with an alpha-value of 0.05. The p-values and gene ontology terms can be found in Supplementary Tables S3-S6. A custom R script was designed to filter redundant GO Terms and plotted in respect to the number of proteins which are differentially regulated (Github: DanielleMStevens/Franco_2020_Proteomics_Paper) (48, 53). Raw data and code can be found in the above repository.

### Phylogenetic analyses

A total of 134 serine protease and 84 peroxidase sequences were obtained from Phytozome using the *C. clementina* v1.0 and *C. sinensis* v1.0 databases (43, 54). Peroxidases were identified using the RedOxiBase Database (http://peroxibase.toulouse.inra.fr/) and the pFAM domain PF00141. Serine proteases were identified using the MEROPs peptidase database (https://www.ebi.ac.uk/merops/index.shtml). Multiple sequence alignments were performed with PRANK (55). A maximum likelihood approach was used to reconstruct phylogenetic trees using RAxML (56). Bootstrapping was performed with 1,000 replicates. The resulting phylogenetic trees were visualized using FigTree (version 1.4.3) (57).

### Activity Based Protein profiling (ABPP)

Leaves from the *C. sinensis* N-33 and Hamlin varieties (n =4 trees) were ground in Tris-HCl pH 8.0 solution. The supernatant was collected and centrifuged at 5,000 x g for 10 min. Citrus extracts were quantified using the Pierce 660 Protein assay kit (ThermoFisher Scientific). The serine protease inhibitor cocktail (50 μM chymotrypsin, 50 μM PMSF, 50 μM of 3,4-dichloroisocoumarin) was pre-incubated for 30 min prior to the addition of the ActivX TAMRA fluorophosphonate (FP) serine hydrolase probe (ThermoFisher Scientific). For the no probe control, 1 μL of dimethyl sulfoxide (DMSO) was added. The serine hydrolase FP probe was added at a final concentration of 2 μM. Serine proteases were labeled for one hour, labeling stopped by adding 2X Laemmli buffer, and the sample incubated at 95 °C for 5 min. Labeled proteins were run on an 8-16% precise Tris-Glycine SDS page gel (ThermoFisher Scientific). Serine protease activity was visualized by in-gel fluorescence using the Typhoon 8600 variable mode Imager (Molecular Dynamics Inc, CA, USA). Significant differences in serine protease activity were detected with a two-tailed unpaired Student’s t-test alpha-value of 0.05.

### Peroxidase gene expression using RT-qPCR

#### Hamlin sweet orange

Total RNA was extracted with the RNeasy Plant Mini Kit (Qiagen) and treated with DNase I (Promega). First-strand cDNA was synthesized from purified RNA with ImProm-II™ Reverse Transcription System (Promega) and diluted 10 times for RT-qPCR to detect related genes with specific primers. Quantitative reverse transcription PCR reactions used the 2xKiCqStart^®^ SYBR^®^ Green qPCR ReadyMix™ (Sigma-Aldrich). PCR consisted of an initial activation step at 95°C for 3 min, followed by 40 cycles of 95°C for 15 s and 60°C for 40 s. All cDNA samples were run in triplicate. The *C. sinensis* GAPDH gene (XM_006476919.3) was used to normalize the expression of each target gene and quantification was performed using the ΔΔCt method. Significant differences (n=4) were detected with a two-tailed unpaired Student’s t-test.

#### N-33 sweet orange variety

Total RNA was extracted using a Trizol (Invitrogen, USA)-based method and treated with DNase I (DNase Max Kit (50), QIAGEN). Quantitative reverse transcription PCR reactions used Power SYBR™ Green RNA-to C_T_™ 1-step kit (Applied Biosystems-Thermo Fisher Scientific, USA). Thermocycling began with a Hold Stage (first step at 48 °C for 30 s, second step 95 °C for 10 min), followed by a PCR stage (first step at 95°C for 15 s, second step 55 °C for 1 min over 40 cycles) in a thermal cycler (Applied Biosystems-QuantStudio 3, Thermo Fisher Scientific, USA). The *C. sinensis* GAPDH gene (XM_006476919.3) was used to normalize the expression of each target gene and quantification was performed using the ΔΔCt method. Significant differences (n=3) were detected with a twotailed unpaired Student’s t-test.

### Peroxidase activity

Peroxidase activity was tested using a previously described protocol (58). Briefly, citrus leaf punches (4 mm diameter) were excised and washed for 1 hr in 1x MS solution with agitation. Each leaf disk was then transferred to a 96 well-microplate containing 50 μL of 1x MS solution. Leaf disks were incubated for 20 h at 25°C with agitation. To observe peroxidase accumulation, leaf disks were removed and 50 μL of 1mg/mL of 5-aminosalicylic acid, pH 6.0, with 0.01% hydrogen peroxide was added and incubated for 1-3 min. The reaction was stopped by adding 20 μL of 2 N NaOH. Optical density readings (OD_600_) were measured using a spectrophotometer. Uninfected samples were set to 100% peroxidase activity and changes in peroxidase activity were calculated based on the ratio of OD_600_ readings (Infected/Uninfected). Significant differences in peroxidase activity were detected with a two-tailed unpaired Student’s t-test.

## Results

### *C*Las induces global changes in the citrus vascular proteome

To investigate changes occurring in citrus tissues infected with *C*Las, we isolated protein from bark samples ten months post-inoculation and performed proteomics analyses. The sweet orange cultivar Washington navel (*Citrus sinensis* (L) Osbeck), which is susceptible to HLB, was graft inoculated or mock inoculated with *C*Las positive budwood. At 10 months postinoculation, *C*Las inoculated navels exhibited severe HLB disease symptoms and detectable *C*Las titers using qPCR (Supplementary Table S7). Total extracted proteins were subjected to tandem mass spectrometry (LC-MS/MS) and MS1 quantification using MaxQuant (41). To increase total protein identifications, the protein database included the *C. sinensis* and *C. clementina* assembled genomes (54, 59). A total of 1,354 citrus proteins were identified using MS1 quantification for infected and uninfected samples (Supplementary Table 8, Figure 1A). Uninfected samples had an average of 1259 protein identifications and infected samples had an average of 1152 protein identifications across all four biological replicates (trees). 329 (24%) proteins were significantly upregulated and 363 (27%) proteins were downregulated in infected samples (Supplementary Table 9 and 10, respectively). Hierarchical clustering based on Euclidean distance revealed that samples clustered according to treatment (Fig. 1A) (45). Similarly, principle component analyses (PCA) differentiated biological samples based on their treatment (Fig. 1B). Taken together, these data indicate that *C*Las infection induces global and reproducible changes in vascular protein expression patterns (Fig 1A-B).

**Fig. 1.**
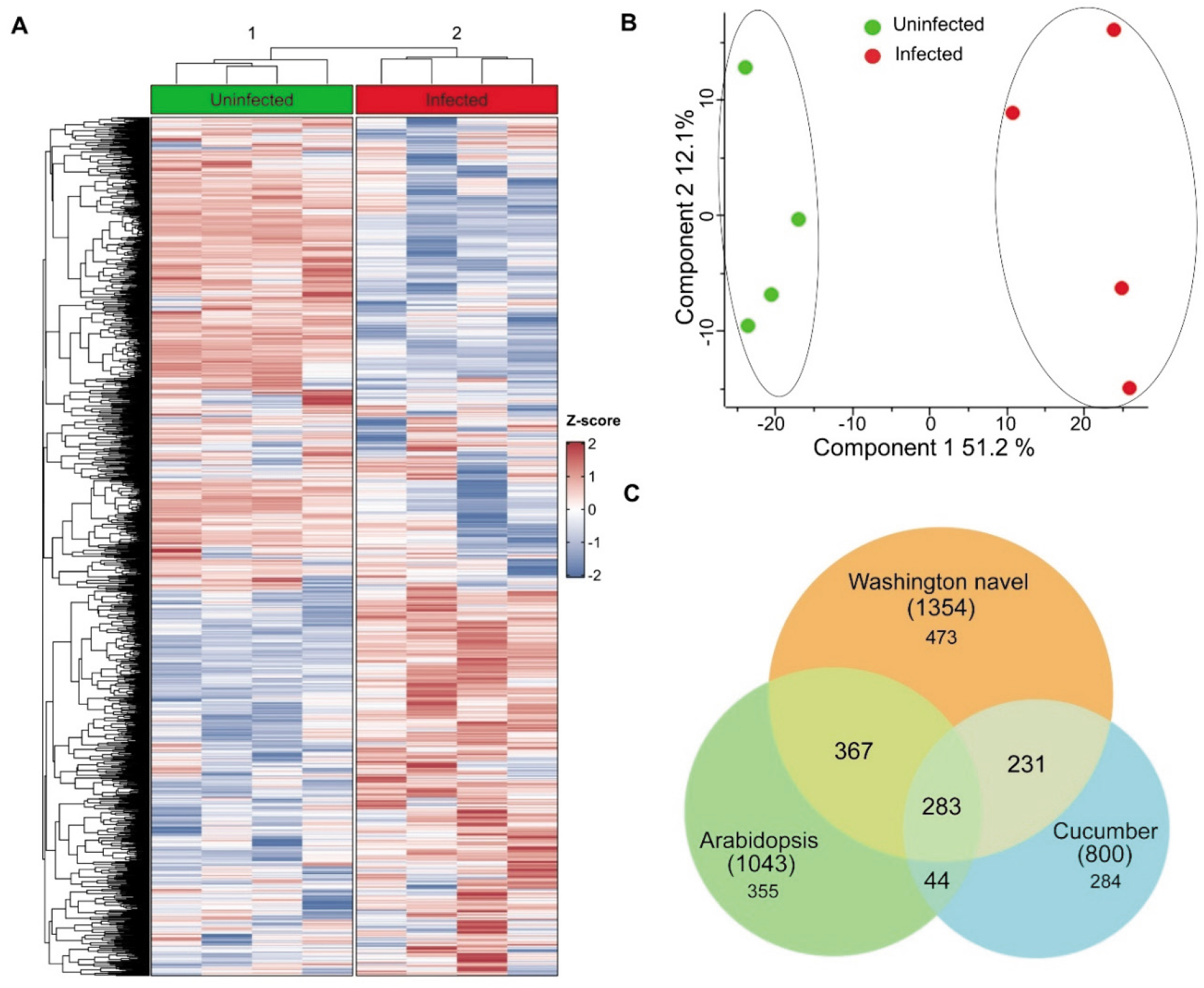
*C*Las induces global changes in protein expression in citrus vascular exudates. **(A)** Heatmap of proteins identified in citrus sap exudates. Vascular proteins were enriched and extracted from mock and *C*Las-graft inoculated Washington navel oranges (*C. sinensis* (L) Osbeck) and subjected to label-free MS1 quantification. Identified proteins are shown in rows, normalized by their mean value, and the Z-score was calculated for hierarchical clustering using Euclidean distance. Columns represent biological replicates (individual trees). Red indicates proteins where expression was increased, and blue indicates decreased protein expression relative to the mean. **(B)** Principal component analyses (PCA) was performed on untransformed label free quantification values in Perseus. **(C)** Comparison of phloem proteins identified in diverse plant species. A total of 1354 proteins from Washington navel identified from this study were compared with previous phloem proteomics studies in *Arabidopsis thaliana* and cucumber (*Cucumis sativus*) phloem proteins using BLAST with an E-value of 1e-10 as a cutoff. The venn diagram indicates protein overlap and was generated using the vennueler R package.

Isolation of pure phloem proteins is extremely difficult and can only be obtained using microneedle puncture or the stylet of a phloem-feeding insect (60, 61). To increase phloem yields, EDTA-facilitated exudation and centrifugation are commonly used (10). However, these approaches are not amenable to citrus exhibiting HLB symptoms due to decreased phloem transport. Therefore, we centrifuged the outer bark to extract vascular exudates and enrich for phloem proteins. We compared the overlap in protein identifications between our study with studies performed in *Cucumis sativus* and *Arabidopsis* (49–51). Citrus homologs for 62% and 64% of the *Arabidopsis* and *Cucumis sativus* proteins were identified, respectively (Fig. 1C). Although these plants represent diverse families (Rutaceae, Brassicaceae and Cucurbitaceae), many of the identified proteins overlapped. These results indicate that vascular proteins were successfully enriched from citrus. However, our samples clearly contain proteins from the phloem as well as tissues surrounding the vasculature. Cellular responses in close proximity to the vasculature can influence the phloem environment and the changes described below likely contain proteomic responses in and surrounding vascular tissue.

To gain insight on the biological and molecular functions altered by *C*Las infection, we performed gene ontology analyses using Phytozome (43). Processes involved in plant growth and development including metabolism, translation and biosynthetic processes were inhibited after *C*Las infection (Fig. 2). For example, many ribosomal proteins and translation elongation factors were downregulated during infection, which can be visualized using the gene ontology enrichment (Fig 2, Supplementary Table 10). Amino acid metabolism was also repressed after infection. Particularly, methionine and glutamate metabolism were significantly dampened after *C*Las infection (Fig 2, Supplementary Table 10). *C*Las has a reduced genome and synthesizes only six of the twenty proteinogenic amino acids *de novo* and must acquire the remaining amino acids from its surroundings (62). HLB susceptible Valencia sweet oranges are more abundant in amino acids like methionine and glutamate in comparison to HLB-tolerant genotypes including curry leaf, orange jasmine and trifoliate orange (63, 64).

**Fig. 2.**
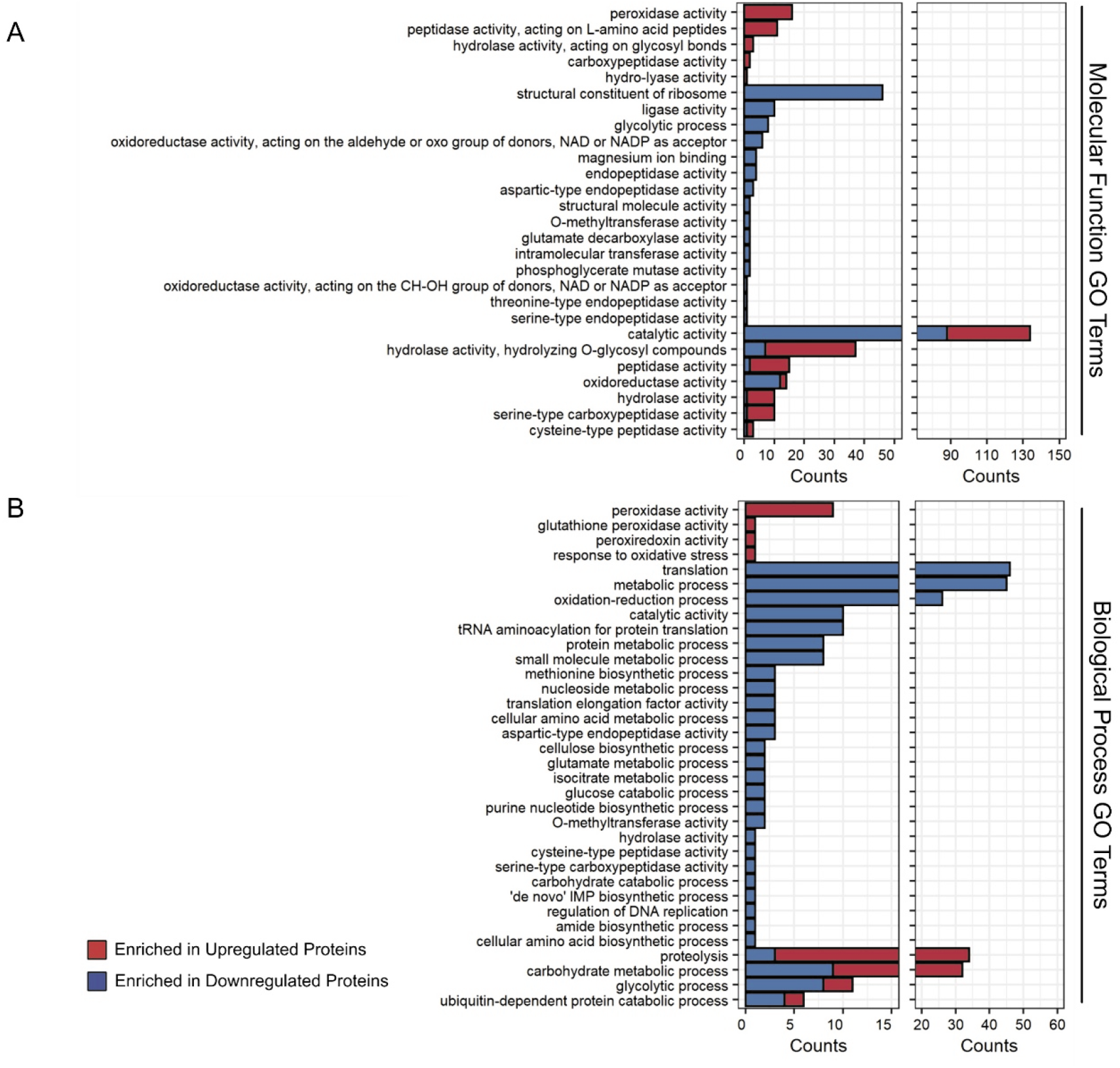
Gene ontology (GO) analyses of citrus proteomics. **(A)** The biological processes and **(B)** molecular function of 692 significantly changing proteins identified by proteomics analyses of enriched vascular proteins from Washington navel ten months post-graft inoculation with *C*Las compared to mock inoculated controls. GO analyses and molecular function was performed using Phytozome 12 (https://phytozome.jgi.doe.gov/). Bonferroni was used for test correction with an FDR rate of <0.05. GO term slimming was performed manually by removing redundant terms used for the same group of proteins. Blue = enriched in downregulated proteins after graft inoculation with *C*Las; Red = enriched in upregulated proteins after graft inoculation with *C*Las.

Proteins involved in pathogen perception were upregulated in infected samples. We identified multiple proteins with receptor-like kinase or receptor-like protein domain architecture that were upregulated during infection, including the receptor-like kinase, Ciclev10030933m, and the LysM domain protein orange1.1g018290m (Supplementary Table 9). Proteins with this domain architecture are frequently implicated in pathogen perception (27, 28, 65). Multiple peroxidases and proteases were more abundant in *C*Las infected samples (Fig 2A, 2B). Both peroxidases and proteases have been established as key components in development and plant defense responses (66–69). Taken together, these results implicate the tradeoff between growth and defense during *C*Las infection.

### Peroxidase transcript and protein abundance increases during *C*Las infection

To gain a greater understanding of citrus peroxidases, we identified the two plant specific peroxidase classes (I and III) in *C. sinensis* and *C. clementina* genomes (70). Class I peroxidases encompass intracellular enzymes such as ascorbate peroxidases and catalases (70–72). Class III peroxidases are a large family with extracellular or vacuolar localization that regulate H_2_O_2_ abundance, oxidation of toxic substrates, and defense responses (69, 73, 74). Using PeroxiBase, 80 peroxidases (13 class I, 67 class III) were identified in *C. sinensis*. For phylogenetic analyses, an additional 4 peroxidases (2 class I, 2 class III) identified from *C. clementina* were included (72) (Fig 3). Maximum-likelihood phylogenetic analyses of these peroxidases were performed using RAxML (56). The identified ascorbate peroxidases cluster appropriately within the class I clade (Fig 3). In the vascular proteome, 22 peroxidases were identified and 10 were upregulated during *C*Las infection (3 class I and 7 class III) (Supplemental Figure 1, Supplementary Table 11). Taken together, these data highlight the abundance of peroxidases in *C. sinensis* and reveal an overall increase in abundance of specific members upon *C*Las infection.

**Fig. 3.**
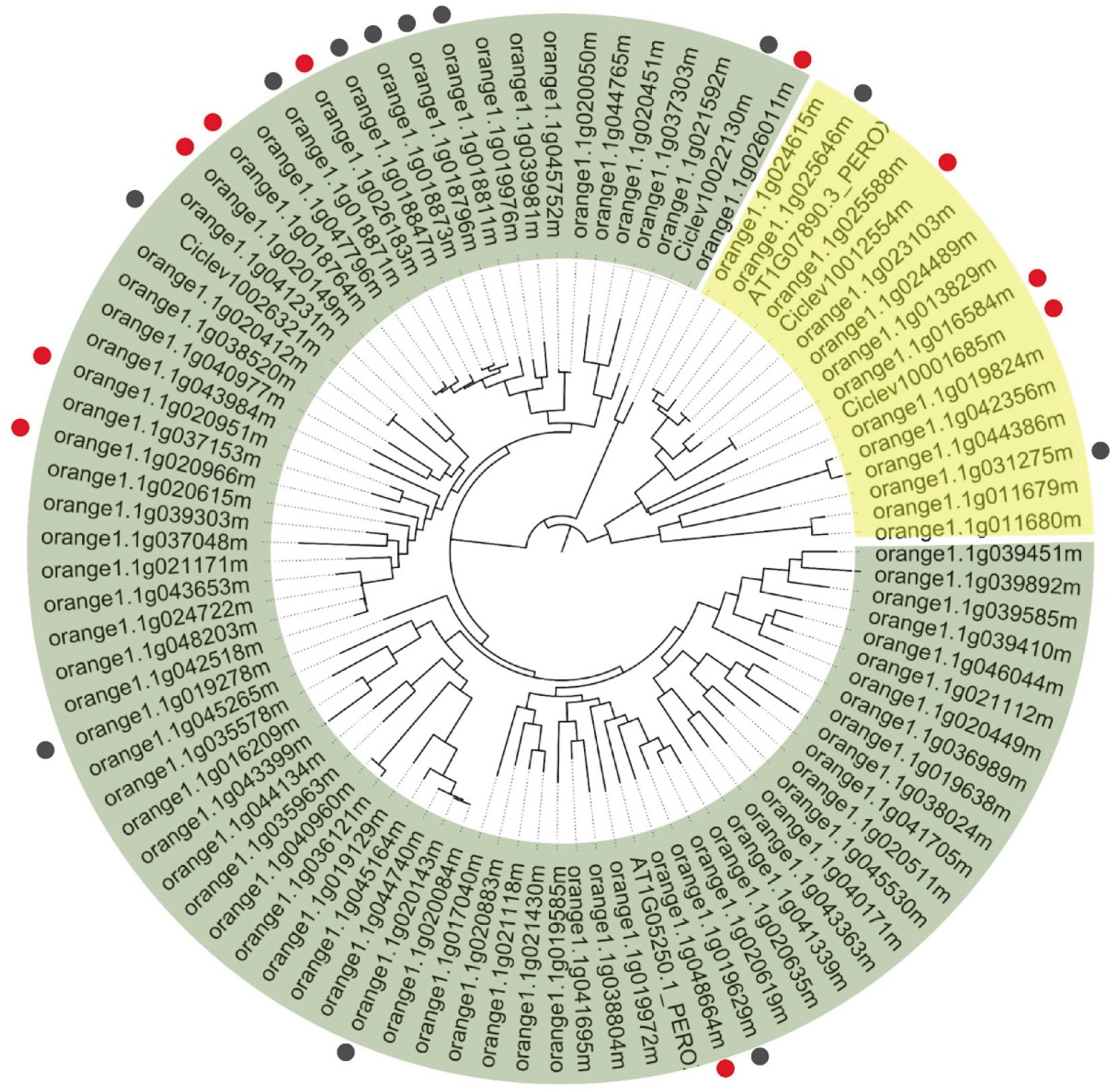
Phylogenetic analyses reveal dynamic changes in specific citrus class I and III peroxidases after inoculation with *C*Las. A total of 15(13, *C. sinensis*, (*Cs*) and 2 *C. clementina*, (*Cc*)) Class I (yellow) and 78 (76 *Cs*, 2 Cc) Class III (green) peroxidases were identified from the citrus genome using the PF00141 Pfam domain and the RedOxiBase database. Two *Arabidopsis thaliana* sequences were used to guide clustering, one class I Peroxidase (AT1G07890.3) and one class III peroxidase (AT1G05250.1). Accession numbers starting with orange1.1g indicate proteins identified from the *C. sinensis* (navel) genome while those starting with Ciclev are those identified from the *C. clementina* (clementine) genome. Redundant sequences from *Citrus clementina* were removed from the phylogeny. A maximum likelihood approach was used to generate the phylogeny with 1000 bootstrap replicates. The resulting phylogeny was visualized using FigTree. A total of 22 peroxidases were identified by mass spectrometry in Washington Navel. Red indicates peroxidases that were detected by mass spectrometry as significantly increased accumulation in *C*Las inoculated Washington navel and grey indicates peroxidases that were detected, but not significantly changing after inoculation.

The abundance of specific enzyme classes does not always correlate with their activity. To investigate if the proteomic changes identified in Washington Navel also occurred in other sweet orange genotypes, we analyzed peroxidase transcript accumulation and enzymatic activity in the *C. sinensis* Hamlin sweet orange variety, which is widely grown in Florida (75). Healthy and symptomatic leaf tissue from Hamlin oranges were sampled one-year after *C*Las transmission by *D. citri* (Fig 4A). To test peroxidase gene expression in Hamlin, qPCR was performed on seven peroxidase genes that were previously identified as induced in our proteomics analyses. Four Hamlin peroxidases were induced in infected samples and three were expressed at the same level in infected and uninfected samples by qPCR (Fig 4B). We investigated peroxidase activity during *C*Las infection using a microplate-based assay to quantify the basal levels of the reactive oxygen species, H_2_O_2_ (58). Overall peroxidase activity decreased in infected Hamlin samples (Fig 4C). The disconnect between peroxidase mRNA expression and enzymatic activity may be due to *C*Las manipulation of enzymatic activity or opposing roles of specific peroxidase enzymes.

**Fig. 4.**
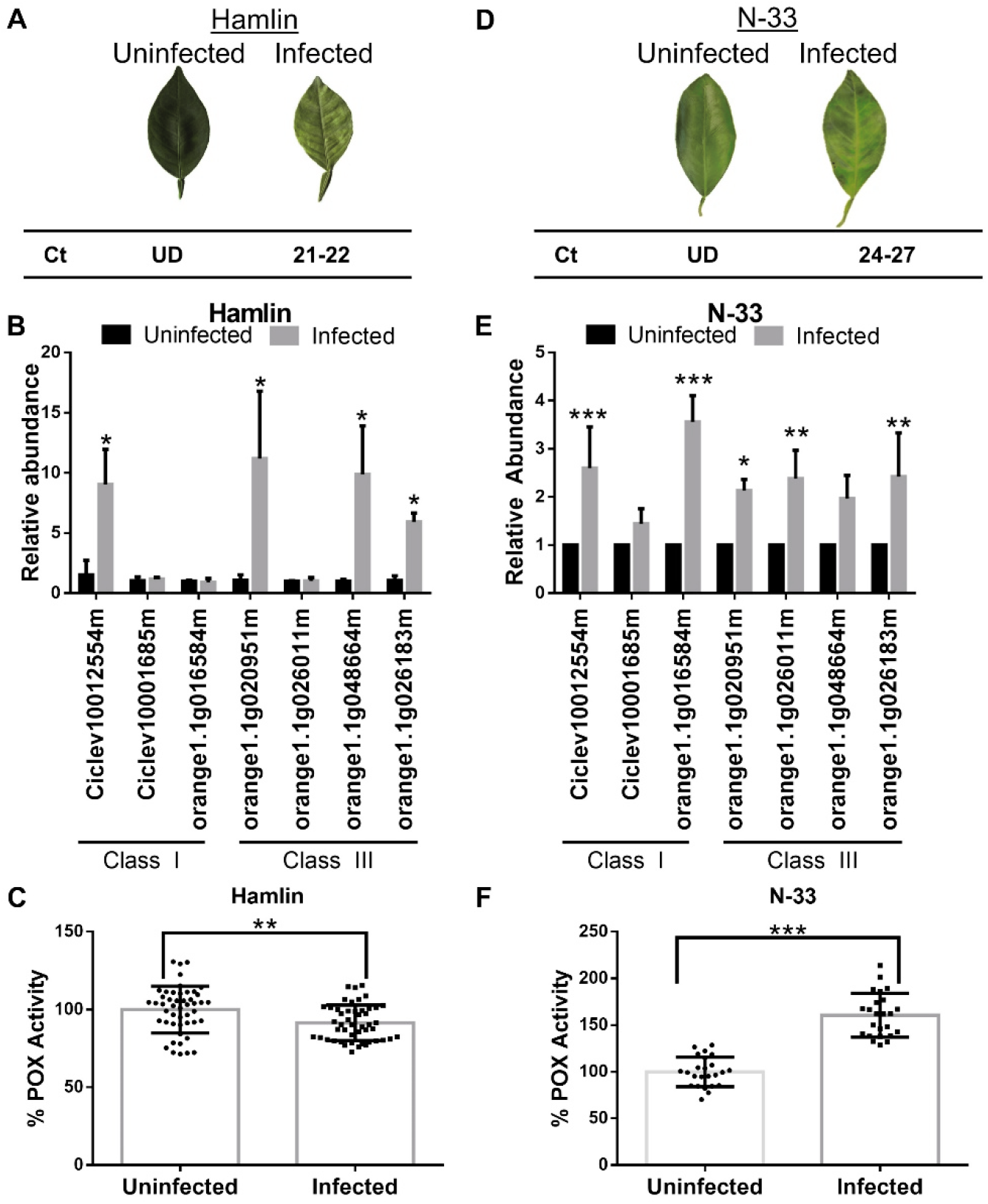
Peroxidase activity and transcript accumulation in two citrus genotypes. Two sweet orange varieties were inoculated with *C*Las using *Diaphorina citri* and analyzed for peroxidase (POX) enzymatic activity and transcript abundance. **(A)** HLB symptoms on representative Hamlin leaves (top) and corresponding Ct value for *C*Las (bottom) one year after feeding with *C*Las positive and negative *D. citri*. Citrus plants were grown under greenhouse conditions. *C*Las titers were detected by qPCR using primers targeting the β-operon (GenBank: AY342001.1). **(B)** Select peroxidase (POX) transcripts increase during infection in Hamlin. Hamlin trees were inoculated as described in (A) and RNA was extracted from the leaves. POXs that were significantly induced by proteomics analyses in Washington navel after *C*Las inoculation were analyzed for transcriptional expression using qPCR. Transcripts were normalized to GAPDH-C (NCBI Rseq: XM_006476919.3). Graphs show means ±SD, n= 4 trees. Significant differences were detected with a two-tailed unpaired Student’s t-test, alpha-value of 0.05. **(C)** POX enzymatic activity decreases during infection in Hamlin. Hamlin trees were inoculated as described in (A) and POX activity was detected on leaf punches surrounding the midrib one-year post-inoculation from the same leaves used for RNA extraction in (B). Graphs show means ±SD, n=12 samples taken from 4 individual trees. Significant differences were detected with a two-tailed unpaired Student’s t-test, p<0.01. **(D)** HLB symptoms on representative N-33 leaves (top) and corresponding Ct value for *C*Las (bottom). Citrus plants were grown in a Texas field where *C*Las was naturally transmitted by *D. citri*. Uninfected leaf samples were taken from plants grown in an adjacent screenhouse. *C*Las titers were detected by qPCR primers targeting the 16s rDNA (GenBank: L22532). **(E)** Select peroxidase transcripts increase during infection in N-33. Leaf samples were taken from trees described in (D) and RNA was extracted from leaf tissue. POXs that were significantly changing in Washington navel proteomic analyses were selected and analyzed as described in (C). Graphs show mean ±SD, n=3 trees. Significant differences were detected with a two-tailed unpaired Student’s t-test, Asterisks represent p values, *=<0.05, **=<0.01, ***=<0.001. Experiments were repeated three times with similar results. **(F)** POX activity increases during infection in N-33. Leaf punches from N-33 leaf samples taken from trees described in (D). Graphs show mean ±SD, n=8 taken from 3 individual trees. Significant differences were detected with a two tailed unpaired Student’s t-test, p<0.001.

N-33 is a Navel orange variety frequently grown in Texas (76). To test peroxidase accumulation under field conditions and with another *C. sinensis* variety, mature N-33 plants were sampled for peroxidase accumulation by qPCR and enzymatic activity. Symptomatic leaves and healthy leaves from an adjacent screenhouse were collected. To validate *C*Las infection, qPCR was performed on seven peroxidases identified in our proteomics analyses as described above (Fig 4D). The transcript abundance of all analyzed peroxidases in N-33 was higher in infected samples, with five demonstrating a statistically significant increase (Fig 4E). Furthermore, peroxidase enzymatic activity was significantly higher in N-33 infected leaf tissue (Fig 4F). While all sweet orange genotypes are susceptible to HLB, mature trees are generally more tolerant than young plants (25, 77). The N-33 genotypes were mature trees and possessed a lower titer of *C*Las compared to the Hamlin genotype and these two factors could influence peroxidase activity. These data highlight the responsiveness of individual peroxidases during *C*Las infection and indicate that the response to *C*Las can be influenced by plant genotype and/or environmental effects.

### Diverse proteases and protease inhibitors dynamically change during infection

Proteolysis is an enriched biological process and molecular function in the mass spectrometry quantification of vascular proteins from *C*Las-infected citrus (Fig 2A, Fig 2B). Plant proteases are divided into seven classes: serine, aspartic, cysteine, asparagine, threonine, glutamate, and metalloproteases playing important regulatory roles in development, reproduction and immunity (78, 79). In the proteome of vascular exudates, both proteases and protease inhibitors were present and dynamically responded to infection. A total of 22 serine, 15 aspartic, 12 cysteine and 2 metalloproteases were identified (Fig 5A).

**Fig. 5.**
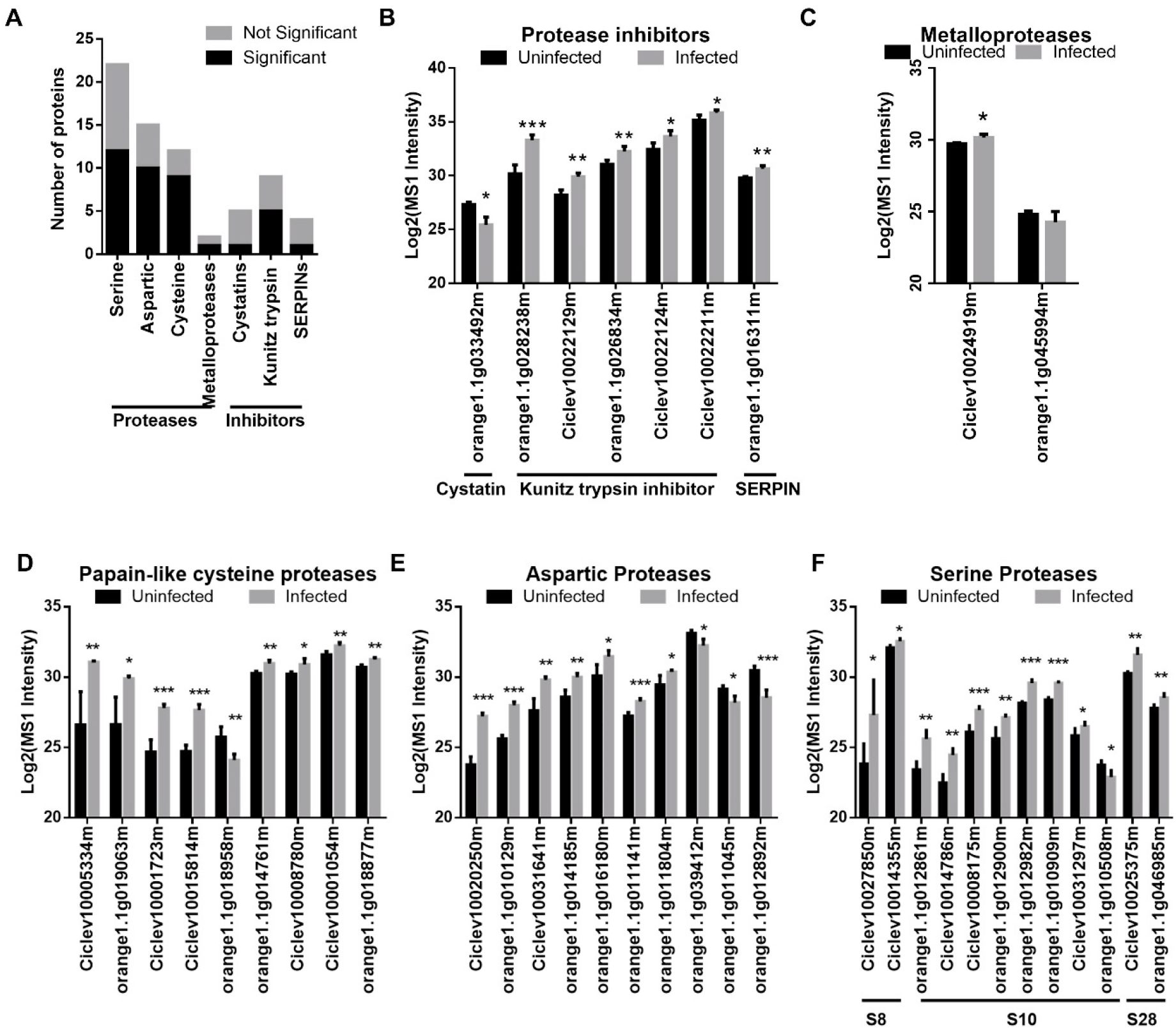
Dynamic interplay between the abundance of proteases and protease inhibitors during *C*Las infection. **(A)** Total number of proteases and protease inhibitors identified by proteomics analyses of enriched vascular proteins from Washington navel ten months post-graft inoculation with *C*Las compared to mock inoculated controls. Four proteases classes comprising: 22 serine proteases, 15 aspartic proteases, 12 cysteine proteases, and 2 metalloproteases were identified. Three protease inhibitor classes were identified: 5 cystatins, 12 serine protease inhibitors (9 Kunitz trypsin inhibitors and 4 SERPINs). Black indicates proteins that were detected by mass spectrometry as significantly increased accumulation in *C*Las inoculated Washington navel and grey indicates proteins that were detected, but not significantly changing after inoculation. **(B-F)** Abundance of specific protease inhibitors and proteases that exhibit differential accumulation by mass spectrometry. Protein abundance is depicted as Log_2_(MS1) intensity. Abundance in mock inoculated is displayed in black and *C*Las inoculated in grey. Graphs indicate means ±SD, n=4 trees. Significant differences were detected by two-tailed t-test with an FDR of <0.5. * = p<0.05, ** p<0.01, *** p<0.001.

Serine proteases are among the largest protease class in plants encompassing over 200 members and are divided into 14 families (79, 80). Members of the subtilases (S8), serine carboxypeptidases (S10), and lysosomal Pro-x carboxypeptidases (S28) families of serine proteases were identified in our proteomics analyses (Fig 5F, Supplementary Fig. 2). Phylogenetic analyses of 67 S8, 48 S10, and 19 S28 serine proteases from *C. sinensis* and *C. clementina* were performed (78). Differentially accumulating proteases did not cluster phylogenetically (Supplementary Fig. 2). S10 proteases generally increased during infection except for one member, orange1.1g010508 (Fig 5F). Subtilisin and serine carboxypeptidases comprise the largest serine protease class, have had prior implications in plant immunity, and are directly inhibited by diverse pathogen classes (79–83). These results indicate that the S8, S10, and S28 serine proteases families are most responsive to *C*Las infection.

Cystatin protease inhibitors target papain like cysteine proteases (PLCPs) and metacaspase-like cysteine proteases with dual roles as defense proteins and regulators of PLCP protein turnover (84). In this proteomic study, five cystatins were identified, but only one decreased in abundance during infection (Fig 5A-B). Diverse serine protease inhibitors were identified in the vasculature (Fig 5A-B). Two potato type I (Pin1) serine protease inhibitors were identified, but did not change during *C*Las infection (Supplementary Table 12). Kunitz trypsin inhibitors and SERine Protease INhibitors (SERPINs) are the most extensively studied and can inhibit diverse protease classes (85, 86). Kunitz trypsin inhibitors are capable of binding to serine, cysteine and aspartic proteases (85). SERPINs are active serine protease inhibitors, but can also inhibit some cysteine proteases (85). Nine Kunitz trypsin inhibitors were identified (five induced) and two SERPINs were identified (one induced) (Fig 5A, B). Kunitz trypsin inhibitors and SERPINs have been demonstrated to have inhibitory effects against pathogens and insects (85). Our proteomics analyses indicate protease activity is tightly regulated during *C*Las infection.

### Serine hydrolase activity dynamically changes during *C*Las infection

To quantify the activity of serine proteases during *C*Las infection, activity-based protein profiling (ABPP) was performed. ABPP enables detection of various enzymes including proteases (87, 88). We used the ABPP probe FP-TAMRA, which depicts the activity of serine hydrolases (80, 89). For ABPP, we used the Hamlin and N-33 varieties grown in the Florida greenhouse and Texas field, respectively. Proteins from four uninfected and four infected samples were extracted from leaf tissue and subjected to ABPP (Fig 6A). A no probe control was included to confirm probe binding to active proteases. Due to a lack of an effective broadspectrum serine protease inhibitor, an inhibitor cocktail containing phenylmethylsulphonyl fluoride (PMSF), 3,4-dichloroisocoumarine and chymostatin was used to verify labeling specificity. In many cases these cocktails are not 100% effective, resulting in some residual protease activity (thus a reduction in signal is indicative of protease activity (80, 89). The inhibitor cocktail was added 30 min prior to the addition of the probe to inactivate serine proteases. Serine protease activity is demonstrated by the presence of a band detected by fluorescence imaging. Active serine protease bands were reduced in the presence of the protease inhibitor cocktail, indicating that ABPP was able to specifically detect serine protease activity (Fig 6B, D).

**Fig. 6.**
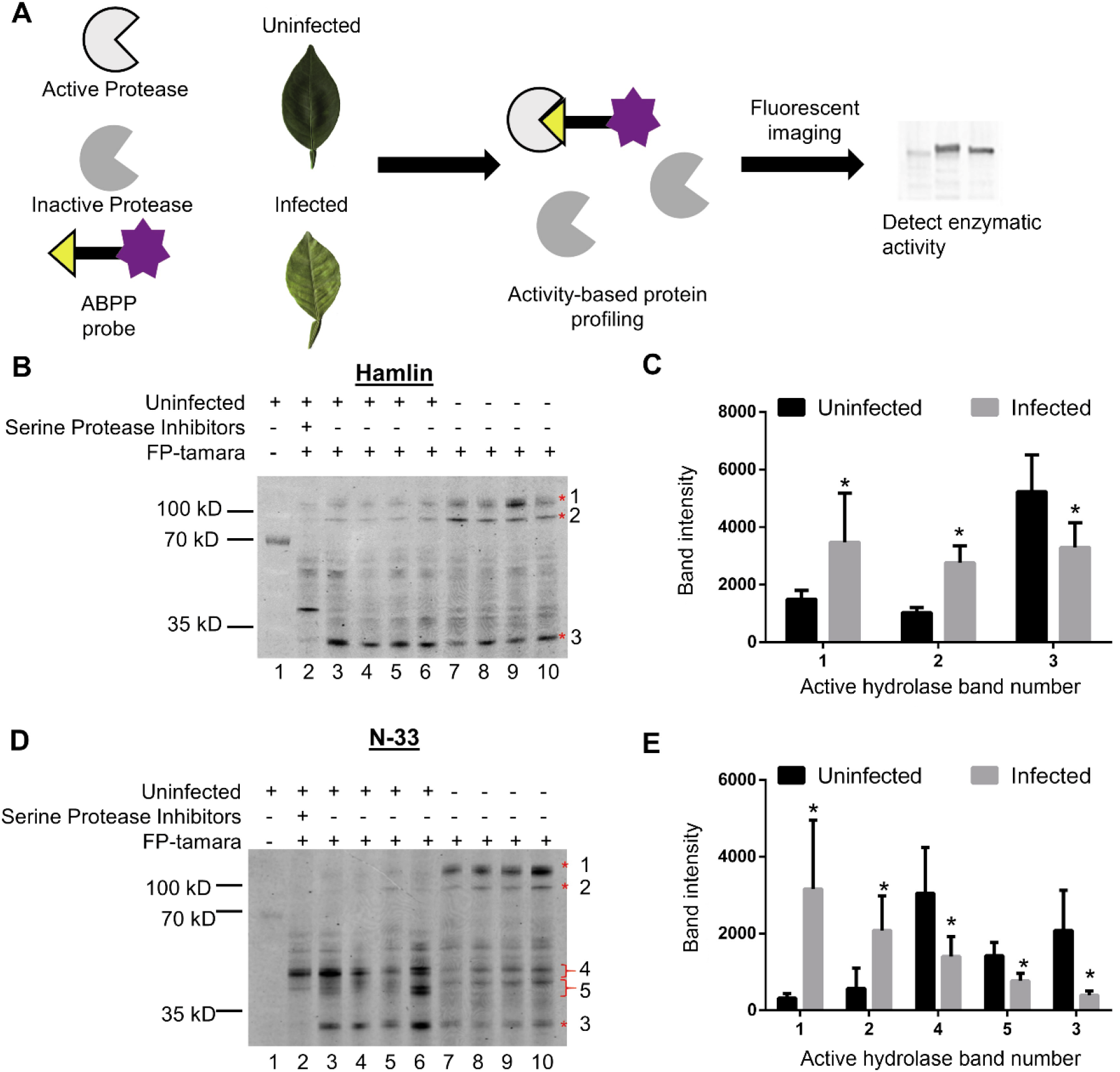
Serine hydrolase activity dynamically changes during *C*Las infection. **(A)** Experimental Strategy. Proteases from uninfected and infected citrus trees were extracted and subjected to activity-based protein profiling (ABPP) using the fluorescently labeled probe, ActivX TAMRA fluorophosphonate (FP) serine hydrolase probe, to capture serine proteases. Visualization of active serine hydrolase bands was performed with the Typhoon 8500 scanner and signals were quantified in Image J. Two sweet orange genotypes were inoculated with *C*Las using *Diaphorina citri* and analyzed for serine protease enzymatic activity. **(B)** ABPP of serine hydrolase activity in leaves of the Hamlin variety one year after feeding with *C*Las positive and negative *D. citri*. Active serine hydrolases were labeled using FP-tamara (lanes 2-10). A serine protease inhibitor cocktail was pre-incubated 30 min prior to the addition of FP-tamara to demonstrate probe binding specificity for serine proteases (lane 2 compared to lane 3). A no probe control was added as a negative control (lane 1, band is the 70kD marker from the protein ladder). Fluorescent signals corresponding to serine protease activity are indicated on the right with an asterisk. **(C)** Quantification of labeled band intensity in (B) using ImageJ. Graphs indicate means ±SD, n=4 trees. Significant differences were detected by two-tailed t-test, * = alpha-value of 0.5. **(D)** ABPP of serine protease activity in leaves of the N-33 variety grown under field conditions. Gel loading is the same as described in (B). Signals corresponding to active serine proteases are indicated by an asterisk. Signals unable to be resolved were grouped together and indicated by brackets. **(E)** Quantification of labeled signal intensity in (C) using ImageJ. Graphs indicate means ±SD, n=4 trees. Significant differences were detected by twotailed t-test, * = alpha-value of 0.05

In Hamlin orange, three distinct protease signals changed during *C*Las infection (Fig 6B). The intensity of bands 1 (~110 kD) and 2 (~100 kD) increased during infection and quantification of the fluorescent signal across four biological replicates demonstrated this increase in activity as statistically significant (Fig 6C). In contrast, band 3 (~30 kD) displayed a significant decrease in protease activity during *C*Las infection. In N-33, bands 1-3 followed the same trend (Fig 6D, E). Two additional groups of serine hydrolase bands were observed in N-33, likely carboxylesterases or other esterases based on their molecular weight (Fig 6D) (89). Due to the inability to resolve individual bands, the average signal was summed in the two different aggregates (bands 4 and 5). Bands 4 and 5 exhibited decreased activity (~40kD) (Fig 6E). These experiments demonstrate the induction and activation of specific serine proteases during *C*Las infection is conserved across sweet orange varieties and diverse environmental conditions.

## Discussion

HLB has caused billions of dollars in economic losses worldwide and continues to threaten the United States citrus industry. To curb HLB losses, a detailed understanding of the mechanisms of disease is essential to develop comprehensive disease management strategies. Various-omics approaches have identified molecular pathways altered during *C*Las infection in leaf, root, stem, and fruit tissues providing a heightened understanding of the responses in diverse tissue types (31, 35, 90–92). For example, proteomics coupled with nutrient profiling demonstrated that *C*Las infection reduces Ca^+2^, Mg^+2^, Fe^+2^, Mn^+4^, Zn^+2^ and Cu^+^ in grapefruit leaves (93). Consistent with these observations, diseased plants supplemented with a cocktail of Ca and Zn resulted in an increase in fruit production (94). However, these studies did not directly address responses within the phloem, the niche where *C*Las resides and propagates. In this study, we quantified proteins in vascular enriched samples to gain a greater understanding of proteomic changes where *C*Las resides.

In Washington navel, we revealed global changes occur in and surrounding vascular tissue, with 51% of the total proteome dynamically changing during *C*Las infection. Upon pathogen perception, cellular resources, which would otherwise be utilized in growth-related processes, are redirected towards defense responses; this is known as the growth-defense tradeoff (95). In this study, we observed evidence for this tradeoff during *C*Las infection, with defense associated proteins induced in infected samples and processes involved in general housekeeping significantly reduced in infected samples. In a parallel study on Washington navel leaf tissue, processes involved in photosynthesis were downregulated while defense related processes were significantly upregulated in *C*Las infected samples (90). These data indicate that the growth-defense tradeoff also occurs in response to vascular pathogens. Washington navel is susceptible to HLB, but clearly responded to *C*Las infection. It is unclear if HLB symptomology is due to a lack of early and robust plant defense, or a result of a slower prolonged autoimmune response, or possibly a combination of the two.

We compared transcription and enzymatic activity of peroxidases and proteases in two genotypes under different environmental conditions. Hamlin sweet orange was grown under controlled greenhouse conditions and mature N-33 sweet oranges were grown in the field. Strikingly, we were able to detect differential transcript expression of peroxidases and differences in enzymatic activity in both peroxidases and proteases in response to HLB infection. Hamlin had fewer peroxidase members differentially changing with a corresponding decrease in POX activity in infected samples. In contrast, N-33 had more peroxidase members with both increased transcript accumulation and POX activity. When assessing serine hydrolase activity, Hamlin similarly presented less drastic changes in activity while N-33 responded more dynamically after *C*Las infection. Although sweet orange varieties are universally susceptible to HLB, multiple studies have suggested mature trees present heightened HLB tolerance than young seedlings (25, 77). These findings are consistent with mature N-33 genotypes exhibiting a greater response to *C*Las.

Reactive oxygen species (ROS) such as superoxide radicals, hydrogen peroxide and singlet oxygen play a central role in plant pathogen defense (96). ROS is produced at the plasma membrane by NAD(P)H oxidases and apoplastic peroxidases (96). ROS also accumulates in different organelles including the chloroplast, mitochondria, and peroxisomes (96). Despite the importance of ROS for defense signaling, an overabundance is cytotoxic and therefore requires tight control (96). In melon, redox-related proteins, including peroxidases, are phloem localized and induced during viral infection (49, 97). In this study, the transcription of seven peroxidases that were more abundant in proteomics analyses, were also upregulated in infected samples from different locations and citrus varieties. ROS regulating enzymes are upregulated at the protein level in grapefruit after *C*Las infection (93, 98). Citrus undergoing thermotherapy to treat HLB exhibited increased peroxidase accumulation (99). Additionally, bioinformatic analyses comparing four RNA-seq studies on HLB illustrated that tolerance is linked with the induction of redox-related genes involved in both young and mature leaf tissues (100). These studies indicate the importance of regulating the opposing roles of redox during *C*Las infection.

Plant proteases are involved in pathogen perception, defense priming, signaling, and programmed cell death (83). For example, proteases release immunogenic peptides from bacteria facilitating perception in host plants (101). To ensure tight control of protease activity, plants evolved protease inhibitors that fine-tune protease activity. Protease inhibitors can also inhibit digestive proteases from herbaceous insects and target pathogens (61, 85, 102, 103). Multiple classes of proteases and protease inhibitors are found in the vasculature of diverse plant species (32, 102–106). In *Arabidopsis*, tomato, and maize, papain-like cysteine proteases (PLCPs) positively regulate immunity and defense hormone accumulation (83, 107, 108). In citrus, PLCPs are localized in phloem-enriched tissue, immune related, and targeted by the conserved *C*Las effector, SDE1 (23). In this study, proteases and their inhibitors were the largest class of dynamically changing proteins during *C*Las infection. In Hamlin and N-33 sweet orange varieties, we observed enhanced and repressed activity of specific citrus serine proteases, suggesting protease enzymatic activity is fine-tuned during *C*Las infection. Whether this inhibition is due to the endogenous activity of plant protease inhibitors, *C*Las inhibition of citrus proteases, or reduced protein accumulation needs further investigation.

Although the serine ABPP probe targets serine proteases, it can capture other enzymatic hydrolases (80, 88, 89). For example, serine hydrolase activity profiling in Arabidopsis apoplastic fluids after infection with the fungal pathogen *Botrytis cinerea* identified a variety of serine carboxypeptidases, prolyl oligopeptidase-like proteins, other hydrolases including carboxylesterases (CXEs) and methylesterases, but very few subtilases (89). Serine ABPP performed in the apoplastic fluids of tomato undergoing programmed cell death after coexpressing the tomato resistance protein Cf-4 and the fungal pathogen *Cladosporium fulvum* effector protein Avr4 identified subtilases (~70kD), serine carboxypeptidases (~50kD), and multiple hydrolases including CXEs and α-hydroxynitril lyases (~40kD) with altered enzymatic activity (80). The banding patterns of sweet orange varieties exhibited bands at ~110kD, ~100 kD, ~40kD, and ~30kD. The significantly changing subtilases in Washington navel are predicted to be ~80kD, and the carboxypeptidases at ~50kD, similar to previously described serine hydrolases (80, 82, 89). It is likely that the larger bands 1 and 2 are subtilases and bands 4 and 5 are carboxypeptidases. These results and prior studies suggest subtilases and serine carboxypeptidases play a large role in mediating plant-pathogen interactions. Future studies identifying the active enzymes after *C*Las infection would provide a profile of active proteases in citrus and future targets to enhance HLB resistance.

Comparative analyses of different citrus genotypes are beginning to reveal the importance of genotype specific responses in defense against *C*Las. RNA-seq analyses between the more HLB-tolerant rough lemon (*Citrus jambhiri*) and susceptible sweet orange (*C. sinensis* (L) Osbeck) demonstrated that the expression amplitude of defense related genes was much higher in the HLB tolerant-rough lemon than HLB-susceptible sweet orange (92). In two susceptible genotypes, Lisbon lemon and Washington navel, the root metabolome and microbiome differed in response to *C*Las. (109). However, both genotypes exhibited a reduction of sugars, the amino acids proline and asparagine, and in some metabolites (109). These important findings are not entirely unexpected as sweet oranges and lemons are different species and were derived from different admixtures (110).

In this study, we observed differences in peroxidase and serine enzymatic activity in two sweet orange varieties, Hamlin and N-33 indicating these enzymatic responses are robust and conserved (109). However, the identity of specific members changed based on experimental or genotype conditions. Future studies investigating diversity of citrus responses to HLB across different citrus genotypes, *C*Las isolates, and environmental conditions will provide a comprehensive understanding of disease development.

## Abbreviations

HLB: huanglongbing;
*C*Las: *Candidatus* Liberibacter asiaticus;
ACP: Asian citrus psyllid;
ABPP: activity-based protein profiling;
PCA: principle component analysis

## Acknowledgements

This research was supported by a USDA National Institute of Food and Agriculture grant #2016-70016-24833 to G.C., V.A., and N.W and grant # 2019-70016-29796 awarded to G.C., and N.W. Additional support was provided by the Citrus Research Board (CRB) project 5200-157. J.Y.F was supported by the USDA National Institute of Food and Agriculture grant #2017-678011-26039. T.W.H.L. was supported by a Rubicon grant of the Netherlands Organization for Scientific Research (NWO).

## Author Contributions

G.C. and J.YF. conceived and designed the study; J.Y.F, S.P.T., Z.P, and F.B.G. performed the experiments. S.P.T., T.W.H.L and D.M.S. performed the bioinformatic analyses. J.Y.F. and G.C. wrote the manuscript with input from all authors.

## Data Availability

Proteomic data are available via ProteomeXchange with identifier PXD020703. Peptide identifications are available via MS-Viewer search key, 9mv75ckfbb.

Gene Ontology analysis pipeline is available via Github: (https://github.com/DanielleMStevens/Franco_2020_Proteomics_Paper)

