## Supplementary figures for "Citrus vascular proteomics highlights the role of peroxidases and serine proteases during Huanglongbing disease progression" (from another version of this preprint: version 2, 2020-04-18)

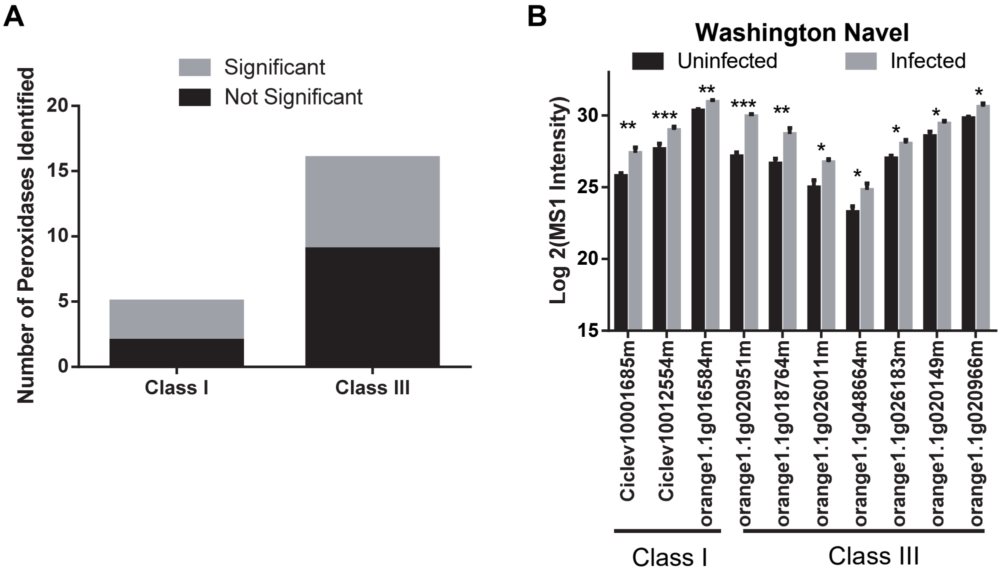


**Supplemental Fig. 1. Class I and class III Peroxidases are induced in the enriched vascular sap proteome. (A)** Total peroxidases identified in the enriched vascular proteome in Washington navel**.** A total of 5 class I peroxidases and 17 class III peroxidases were identified. Grey indicates significantly changing proteases and black indicated identified peroxidases. **(B)** Peroxidase protein abundance is induced in infected Washington Navel. Proteins from mock-inoculated and *C*Las-inoculated plants sampled ten months post *C*Las inoculation from the vascular proteomics study were subjected to MS1 quantification. MS1 intensity was normalized using log2 transformation (y-axis). A total of 10 significantly changing peroxidases were identified, all of which were significantly induced. Graphs indicate means ±SD, n=4 individual trees. Significant differences were detected by two-tailed t-test, * = $p<0.05$, **=$p<0.01$, ***= $p<0.001$.

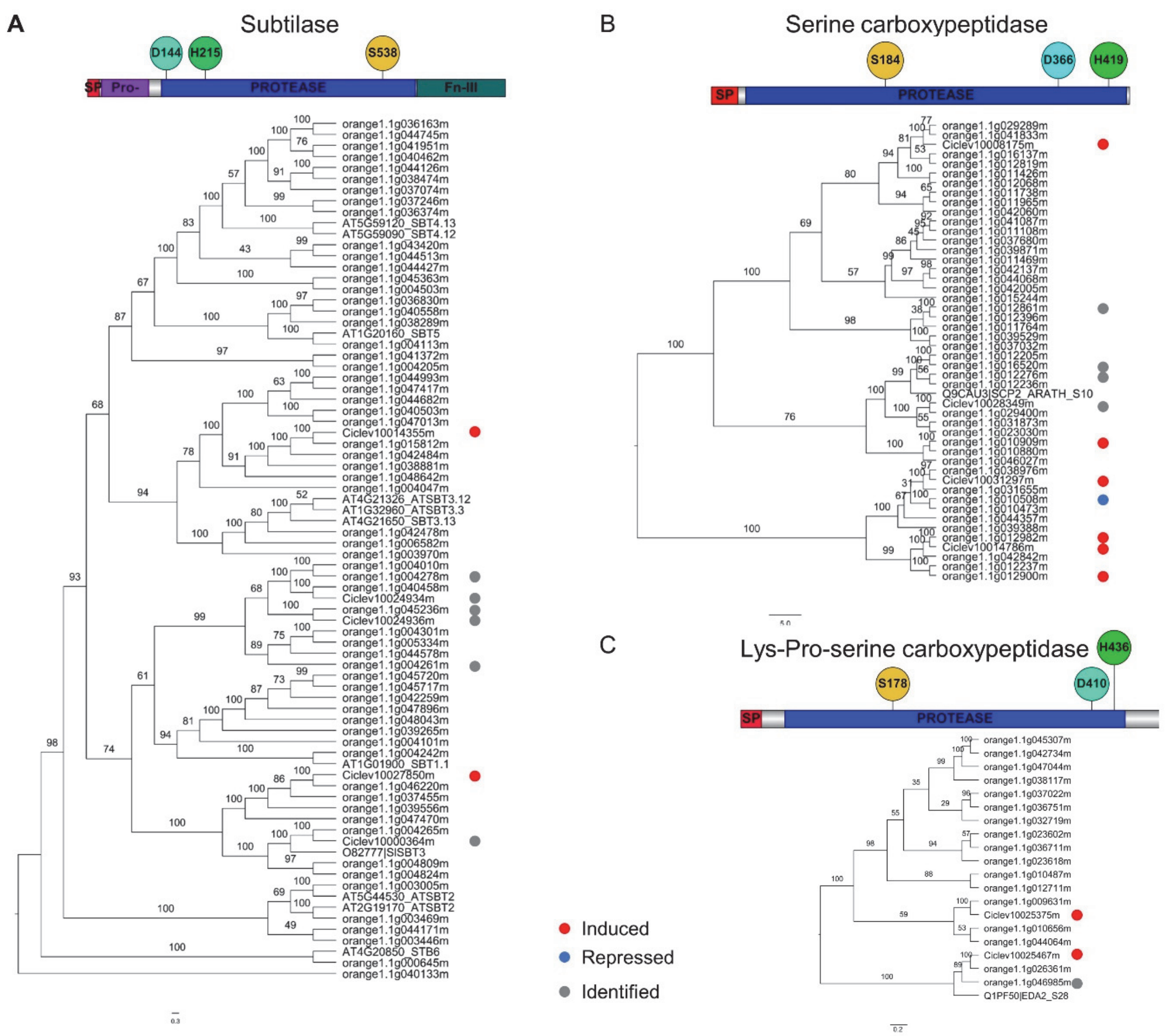


**Supplemental Fig. 2. Phylogenetic analyses of serine proteases identified in Washington Navel during *C*Las infection.** Serine proteases were mined from the *Citrus sinensis* and *C. clementina* genomes and subjected to phylogenetic analyses. Accession numbers starting with orange1.1g indicate *C. sinensis* proteins while those starting with Ciclev indicate *C. clementina* proteins. Redundant sequences from *C. clementina* were removed from the phylogeny. A maximum likelihood approach was used to generate the phylogeny with 1000 bootstrap replicates. Red = induced, blue = reduced, grey = identified in vascular proteomics. *Arabidopsis thaliana* (AT) or *Solanum esculentum* (sp|O82777) sequences were used to guide clustering. **(A)** Subtilase domain architecture and phylogeny. The tomato SBT3 (sp|O82777) was used as a representative subtilase. Subtilases contain a signal peptide (red), an autoinhibitory domain (purple), a protease domain (blue) and may contain a fibrolectin III (FnIII) domain (green). Catalytic residues are indicated. **(B)** Serine carboxypeptidase domain architecture and phylogeny. The *A. thaliana* serine carboxypeptidase 2 (SCP2, AT1G73300) was used as a representative for the domain architecture. **(C)** Lys-Pro-serine carboxypeptidases domain architecture and phylogeny**.** The *A. thaliana* EDA2 (AT2G18080) was used as a representative for the domain architecture.
